# Focus-tunable microscope for imaging small neuronal processes in freely moving animals

**DOI:** 10.1101/2020.10.31.363507

**Authors:** Arutyun Bagramyan, Loïc Tabourin, Ali Rastqar, Narges Karimi, Frédéric Bretzner, Tigran Galstian

## Abstract

Miniature 1-photon microscopes have been widely used to image neuronal assemblies in the brain of freely moving animals over the last decade. However, these systems have important limitations for imaging in-depth fine neuronal structures. We present a novel subcellular imaging 1-photon device that uses an electrically tunable liquid crystal lens to enable a motion-free depth scan in the search of such structures. Our miniaturized microscope is compact (10 mm × 17 mm × 12 mm), lightweight (≈ 1.4 g), provides fast acquisition rate (30-50 frames/second), high magnification (8.7x) and high resolution (1.4 μm) that allow imaging of calcium activity of fine neuronal processes in deep brain regions during a wide range of behavioural tasks of freely moving mice.

## Introduction

Over the last decade, the development of miniaturized microscopes has been instrumental in studying neural circuits of deep brain regions in freely behaving animals (Aharoni and Hoogland, 2019; Miyamoto and Murayama, 2016; Yang and Yuste, 2017; Zong and Chen, 2019). Although it is well established that neurons do not function as simple point processes, less is known about dendritic and synaptic integration in neural circuit function. Indeed, we have a rather good knowledge of how neuronal assemblies contribute to specific behaviors, nevertheless, little is known about the spatiotemporal dynamics of dendrites and dendritic spines, wherein the information is integrated and processed. Therefore, imaging small neuronal processes in freely behaving animals using genetically encoded calcium indicators or voltage-sensitive dyes (Greotti and De Stefani, 2020; Mollinedo-Gajate et al., 2019; Oh et al., 2019) appears to be the next step toward the understanding of brain functions in normal and neuropathological conditions.

Benchtop 2-photon microscopy has been the first approach to record calcium transients from spines, dendrites, and fine processes in the brain of head-restrained animals (Li et al., 2018; Mittmann et al., 2011; Voigts and Harnett, 2018). Although promising, head-restrained models cannot address normal behaviors such as grooming, locomotor gaits, spatial navigation, or social interactions that require free movements of animals. Furthermore, head fixation causes aberrant sensory feedback on the neck and the head, leading to stress, fatigue, and consequently abnormal neuronal processing conditions that are no longer representative of normal functioning (Aghajan et al., 2015; Brand et al., 2012; Ravassard et al., 2013; Sławińska et al., 2012; Thurley and Ayaz, 2017).

To overcome this limitation, portable miniaturized high-resolution devices have been developed (Helmchen et al., 2013, 2001; Ozbay et al., 2018, 2015; Piyawattanametha et al., 2009; Rivera et al., 2011; Zhang et al., 2012; Zong et al., 2017). Among these, a variety of fiber scanning techniques have demonstrated the capacity to record different small neuronal structures (Helmchen et al., 2013, 2001; Ozbay et al., 2018; Rivera et al., 2011; Zhang et al., 2012). However, these systems have multiple drawbacks regarding their size, weight, complexity of movement correction algorithms, and the sampling rate (≤ 5 frames/second (fps)) that limit experiments primarily to structural imaging (Mano et al., 2019). Recently, a miniaturized 2-photon system has reported the use of microelectromechanical systems that has enabled a higher acquisition rate (≈ 40 fps for 256×256 pixels^2^) for Ca^2+^ imaging of small dendritic trees and spines (Zong et al., 2017). Although suitable for functional studies, this system is complex to build and requires the use of hardly accessible dispersion-adapted optical fibers (Zong et al., 2017). Miniaturized systems with external microelectromechanical systems are less complex (Ozbay et al., 2018, 2015), but still require thick fiber bundles that significantly decrease the optical resolution and degrade the quality of images. Finally, miniaturised multiphoton systems use lasers that are bulky, expensive, and barely compatible with the development of future wireless devices. The use of large-diameter optical probes (≥ 1.4 mm) here is also problematic, as it requires tissue ablation above the region of interest prior the implantation of probes in the brain. This can damage the brain, alter neuronal activity, and require multiple series of control experiments (Lee et al., 2016).

Consequently, there is a lack of minimally invasive, accessible, and miniaturized systems adapted for the functional imaging of fine neuronal processes at different depths in the brain of freely behaving animals. To address these issues, we have developed a novel tunable liquid crystal lens (TLCL) that enables electrical shift of the imaging plane within a homemade miniaturized 1-photon system. Our system is lightweight (≈ 1.4 g), has a high spatiotemporal resolution (1.4 ± 0.1 μm) and electrical depth scanning capacity of ≈ 98 μm that overpasses existing systems (Cai et al., 2016; Ghosh et al., 2011; Liberti et al., 2017) and allows us to image Ca^2+^ dynamics of small neuronal processes in the cortex of freely moving mice over a wide range of behaviours. We think that this approach could enable the development of new tools to investigate the dynamics of dendrites and dendritic spines, thus broadening our understanding of synaptic integration and processing in physiological and pathological conditions in freely behaving animals.

## Results

### Key features of the developed device

To overcome the existing depth scanning limitation and to build the most efficient miniaturized system possible, different optical, mechanical and electrical criteria were considered. The 1-photon approach was privileged over the 2-photon to design the most accessible and wireless-friendly system possible. We used a CMOS sensor with high sampling rate of 30-50 fps to enable functional imaging of Ca^2+^ dynamics within neurons and their processes. To image and to record from small neuronal structures, we built a high-magnification system (Figure 1) inspired from a recently reported miniaturized 1-photon microscope (Bagramyan, 2020). The resulting pixel/object ratio of ≈ 2.9 allowed us to distinguish small dendrites, average size ≈ 1 μm spines (Arellano et al., 2007) in a 150±10 μm field of view. To further facilitate the imaging of small structures, we built a special gradient index (GRIN) probe assembly with low optical aberrations compared with those of conventional singlet probes (Bagramyan and Galstian, 2019; Lee and Yun, 2011) (Figure 1). To image neuronal activity in three dimensions, we developed and integrated a TLCL that allowed a motionless electrical scanning through a ≈ 98 μm depth. To diminish tissue damage, our tunable device was optically coupled to a minimally invasive, small-diameter (0.5 mm) GRIN probe assembly (Figure 2D). Finally, our epifluorescence configuration allowed us to image and record calcium activity from fluorescent proteins and calcium indicators, such as GCaMPs.

**Figure 1.**
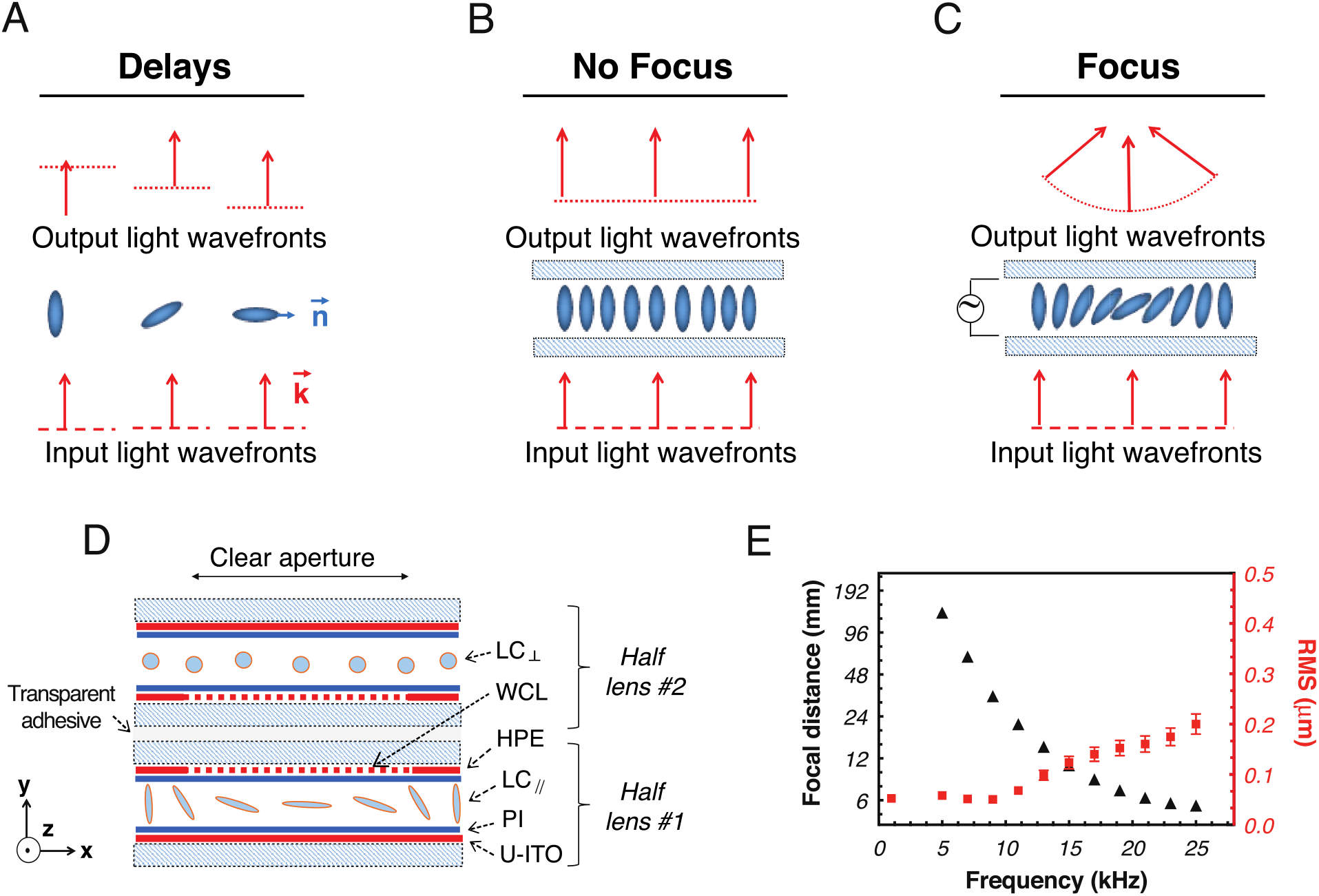
Liquid crystals. **(A-B)** Schematic demonstration of motionless focusing by reorientation of liquid crystal molecules (filled ellipses) **(A)** Different phase delays for different angles between the light wavevector **k** and the liquid crystal optical axis **n**, **(B)** uniform molecular orientation generates uniform phase delay; **(C)** non-uniform molecular orientation generates spherical phase delay and light focusing. **(D)** Schematic demonstration of the structure of the TLCL used (see the main text for details); WCL: weakly conductive layer, PI: polyimide alignment layer, HPE: hole patterned electrode, U-ITO: uniform ITO electrode; **(E)** Control of the focal distance of the TLCL by the frequency of the excitation electrical signal (AC square shaped, at 3V_RMS_) and corresponding RMS aberrations. Error bars for focal distance (7%, not visible) and RMS error (10%) correspond to the maximal measurement fluctuation of the Schack-Hartmann detector.

**Figure 2.**
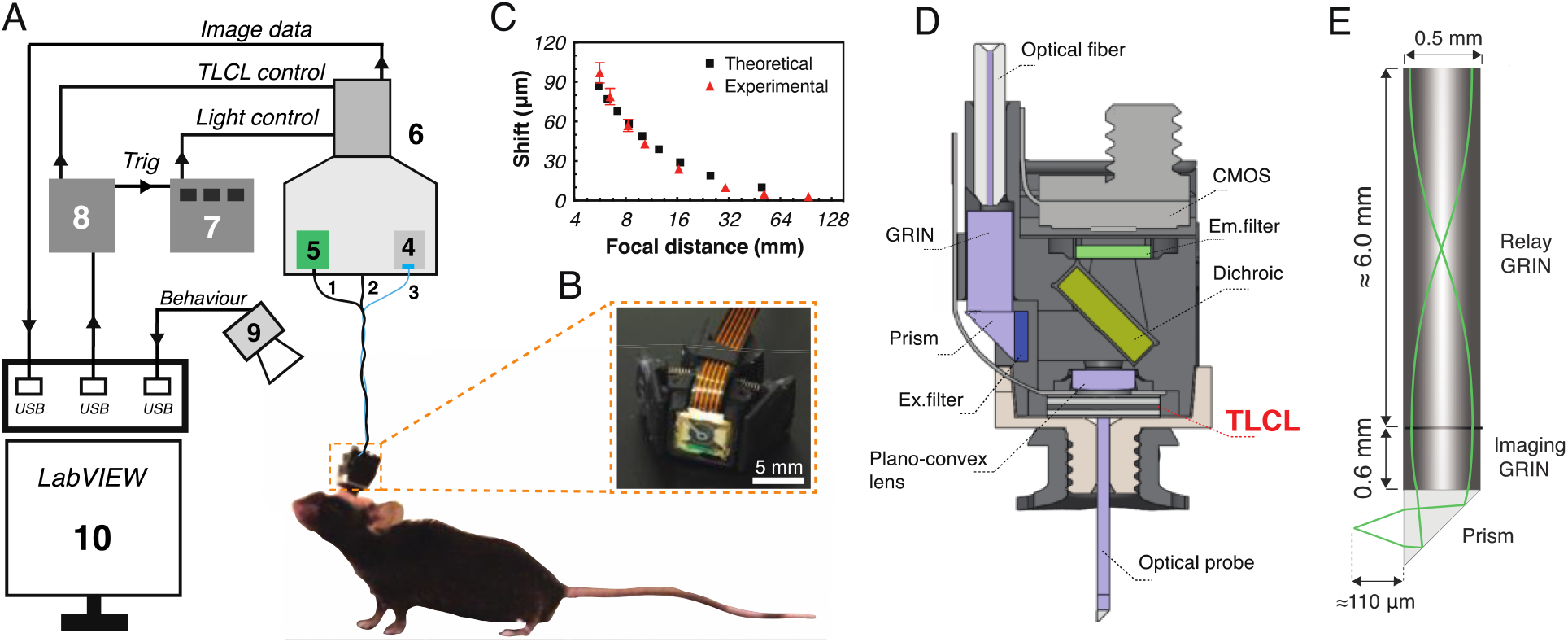
Schematic presentation of the ensemble of the imaging system. **(A)** Schematic of the system: 1. Extension cable for the CMOS; 2. TLCL wires; 3. Optical fiber; 4. Light source; 5. DAQ_CMOS_; 6. Rotary joint system; 7. LED driver; 8. DAQ_LabVIEW_; 9. Camera to record behavior; 10. LabView user interface. **(B)** Photo of the TLCL within the mDS1s. **(C)** Electrically induced focal shift of the mDS1s simulated using Zemax optical software. Experimental results were obtained by measuring the full width at half maximum of the point spread function of 1 μm diameter fluorescent (online Methods). Experimental shift error corresponds to the standard deviation (n=3). **(D)** Schematic representation of optical components. Light from the optical fiber is first collimated by a large-diameter GRIN lens and then reflected by the 45° prism. The excitation filter is used to preserve the desired wavelengths and the dichroic mirror to reflect light into the plano-convex lens, towards the TLCL. Excitation light is then guided within the optical probe assembly to finally reach the object/structure of interest. The emission (fluorescent) light passes back through the probe to reach first the TLCL, then the plano-convex lens. The latter focuses the image on the CMOS sensor passing first through the dichroic mirror and the emission filter. **(E)** Schematic side view of the optical probe with 3 components: 45° prism, imaging (NA = 0.42) and rod GRIN (NA = 0.2) lenses.

### Design of the tunable liquid crystal lens

Three mechanisms enable depth scanning: stepper motor mechanical movement, mechanical deformation (e.g., bending the interface between two liquids), and motionless changes of the focal distance of the lens by changing the refractive index of the lens material (Galstian, 2013). Mechanical movement is well-known, but the components are bulky, heavy, and fragile (Grewe et al., 2011). Interface bending requires high voltage, provides less focus tuning and is limited in the choice of aperture for the lens. Indeed, the smallest available electrowetting lens has a diameter of 1.6 mm. By comparison, motion-free focusing by changing the refractive index of a liquid crystal lens has none of these limitations. Electrically variable focusing with a liquid crystal lens is achieved by taking advantage of the following properties of the liquid crystal lens: light will propagate with different phase delays depending upon the angle between its incident wave vector **k** and the average molecular orientation (the optical axis) of the liquid crystal, often represented by a unit vector **n** (Figure 1A) (Gennes and Prost, 1993).

Most importantly, one can easily reorient the optical axis **n** by applying very small electric field. Hence, liquid crystal molecules can be aligned in a non-uniform way (Figure 1C) to generate a spherical delay in the phase of transmitted light. This is the key to electrically variable focusing (Galstian et al., 2017, 2016). Gradual recovery of uniform alignment of the liquid crystal molecules (Figure 1B) can be used to adjust the focusing effect.

Liquid crystal optical devices have numerous advantages such as low electrical power consumption, high optical power, low weight and small size (Galstian, 2013). Although this technology has been successfully introduced in a variety of consumer electronic products (e.g., DVD readers, webcams, mobile phones), its integration in neuroscience imaging applications has been limited to the macroscopic system (Bagramyan et al., 2017). We have designed and built (Figure 1D) a small (5 mm × 5 mm × 0.5 mm), lightweight (≈ 0.1 g), low power consumption (≈ μW) TLCL that is perfectly suited for miniaturized neuroimaging systems (Ghosh et al., 2011). Moreover, the small 0.5mm clear aperture of our TLCL enables the direct coupling with small diameter optical probes (Figure 2E) and the high (≈ 180 dioptres) optical power of our lens provides much higher depth-scanning ability (x9) compared to current commercial devices (Ozbay et al., 2015). The focal distance of the TLCL can be controlled using the voltage or/and the frequency of the electrical excitation (Figure 1E, Figure 1-figure supplement 1)(Galstian et al., 2017).

### Optical probe

Our lens assembly was well suited for imaging deep brain regions. The probe consisted of the three optical elements presented in Figure 2E. First, we used a 45-degree prism with a sharp tip end that facilitated the insertion of the optical probe into the brain. The prism also allowed us to image from the side rather from the front (standard flat tip) where the tissue tends to compress due to the insertion of the probe. Second, we choose a high numerical aperture imaging GRIN to maximize fluorescence collection (Figure 2E). Finally, to transmit the collected light to the rest of the optical system we used a GRIN rod lens with low optical aberrations (Lee and Yun, 2011).

### System schematic

To drive the TLCL, two fine wires (Figure 2A, 2) were conducted from the micro-endoscope to the custom made rotary joint system (Figure 2A, 6) that prevented the twisting of the cable and allowed to preserve the natural movement of the animal (Figure 2-figure supplement 1). To generate an AC signal in the working frequency range (1-30 kHz), a data acquisition system (Figure 2A, 8) with a high sampling rate (up to 900 kS/s) was chosen. Finally, LabView software was used for the dynamic control of the TLCL and triggering of 4, 5, 9 (Figure 2A) for time-lapse imaging over long periods of time.

### Optical characterisation of the assembly

Before assembling the micro-endoscope, we characterized the optical performance of the TLCL. A low voltage square-shaped AC signal was used to change the focal distance (Figure 1E). The TLCL achieved a minimal focal distance of 5.6 mm while maintaining very low, 0.2 μm RMS, aberrations (Figure 1E).

Prior the assembling, the experimentally obtained optical performances of the TLCL were used in Zemax optical studio software to simulate the focal adjustment capability of the final device (Figure 2C). The experimental focal shift of the assembly was measured (online Methods) using fluorescence microspheres (diameter ≈ 1.0 μm) mounted with a thin (≈ 4 μm) cover glass that provided the ability to scan through the entire working distance of the probe (≈ 110 μm). The experimental results agreed with the theoretical predictions (Figure 2C).

We then characterized the changes in optical resolution during electrical shift of the working distance of the final assembly (Figure 2D) by measuring the point spread function of fluorescent microspheres (online Methods). The results in Figure 3A show a slight decrease in resolution due to an increase in optical aberrations (Figure 1E) and light scattering by the TLCL. Average values of lateral and axial resolutions were, respectively, 1.4 ± 0.1 μm and 15 ± 2 μm. Intensity profiles in Figure 3E show that neurons could be clearly identified through the entire range of depth shift with the TLCL.

**Figure 3.**
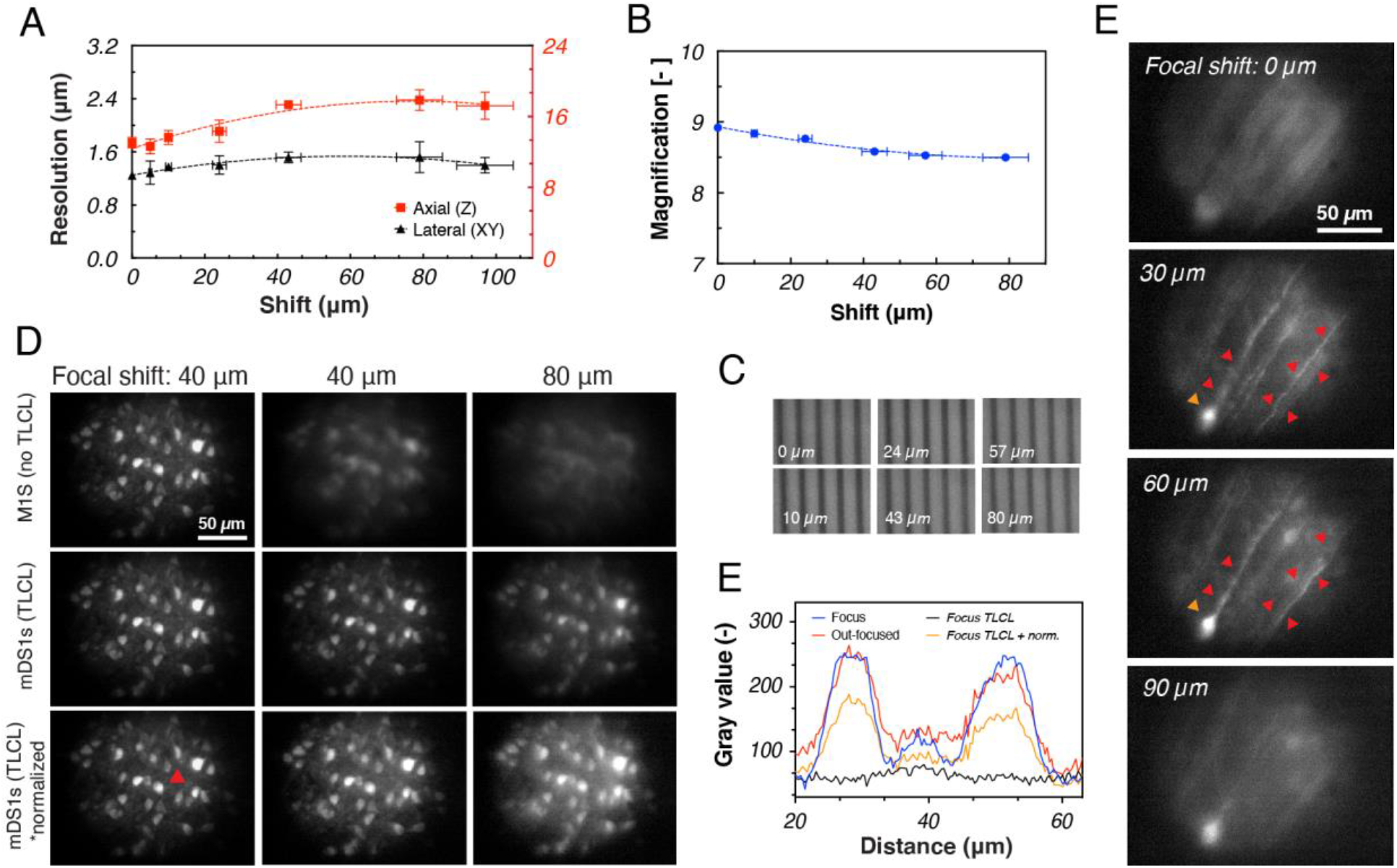
Electrical focal shift and characterisation of optical parameters of the mDS1s. **(A)** Resolution and **(B)** magnification during electrical focal shift. Doted lines correspond to the second order polynomial fits. Resolution error correspond to the standard deviation (n=3) and magnification error is ≈ 1% (n=4). **(C)** Scale bar samples imaged at different depths using the TLCL. Spacing between the lines = 10 μm. **(D)** *In vitro* depth imaging of GCaMP6 expressing neurons. The 1^st^ row presents a sequence of images obtained with a miniaturized 1-photon system (M1S) with a fixed imaging plane. The blur within the mechanically out-focused images (2^nd^ and 3^rd^ columns) cannot be compensated. The 2^nd^ row presents a sequence of images obtained with our miniaturized depth scanning 1-photon system (mDS1s). The electrical depth adjustment allowed to re-focus the images and to reveal the presence of neurons. The 3^rd^ row presents intensity normalization of the 2^nd^ row based on the fluorescence level of the central neuron (indicated with the red arrow). The thickness of slice is 100 ± 10 μm. **(E)** Intensity profile through the dotted line of interest in **D** (3^rd^ row, low-left corner) for the 80 μm focal shift. **(F)** The electrical scan shows various neuronal structures at different depths (indicated in the top left) within the *in vitro* slices presented in **A.**

Having confirmed that our device is suitable for adaptive focusing, we then imaged different brain structures *in vitro*. As demonstrated in Figure 3F, the electrical scan, made with the TLCL, allowed us to image through different layers of fixed tissue preparations. The blurry sections within the first (shift = 0 μm) and last (shift = 90 μm) images were brought back into focus (shift = 30 μm and 60 μm) to clearly show the presence of various dendritic structures (Figure 3F).

### Imaging of dendrites and spines activity in animals

We designed our miniaturized 1-photon device to image fine neuronal processes that are fundamental to the functioning of the brain. Using our system, we performed time-lapse imaging of calcium transients in small neuronal processes of freely moving mice. TLCL was used to adjust the position of the focal plane at the optimal imaging depth where multiple fine neuronal processes were in focus. We then recorded calcium activity (Figure 4-video supplement 1) while the animal was exploring its new cage area (Figure 4A, B). Data were analyzed to investigate the correlation between the activation patterns of different neuronal structures (Figure 4B). The Ca^2+^ traces of a few neurons (Figure 4B, ROIs: 12, 23, 31) showed a very low correlation (Figure 4D) with the rest of the structures, suggesting that these might be a specific population of cells with different activation patterns. Imaging of finer structures (Fig. 4C) revealed high correlation between the activities of dendritic branches **B2** and **B1** (Figure 4D), suggesting that they belong to the same neuron (Figure 4D, ROI: 1). However, the activity of near-passing branch **B3** (morphologically similarity to B2) had a higher correlation with **B4** rather than **B1.** Thus, the analyses of calcium traces (Figure 4C) allowed us to correlate the activation patterns of different fine neuronal structures (Figure 4D), in addition to documenting their structural profiles (Figure 4E).

**Figure 4.**
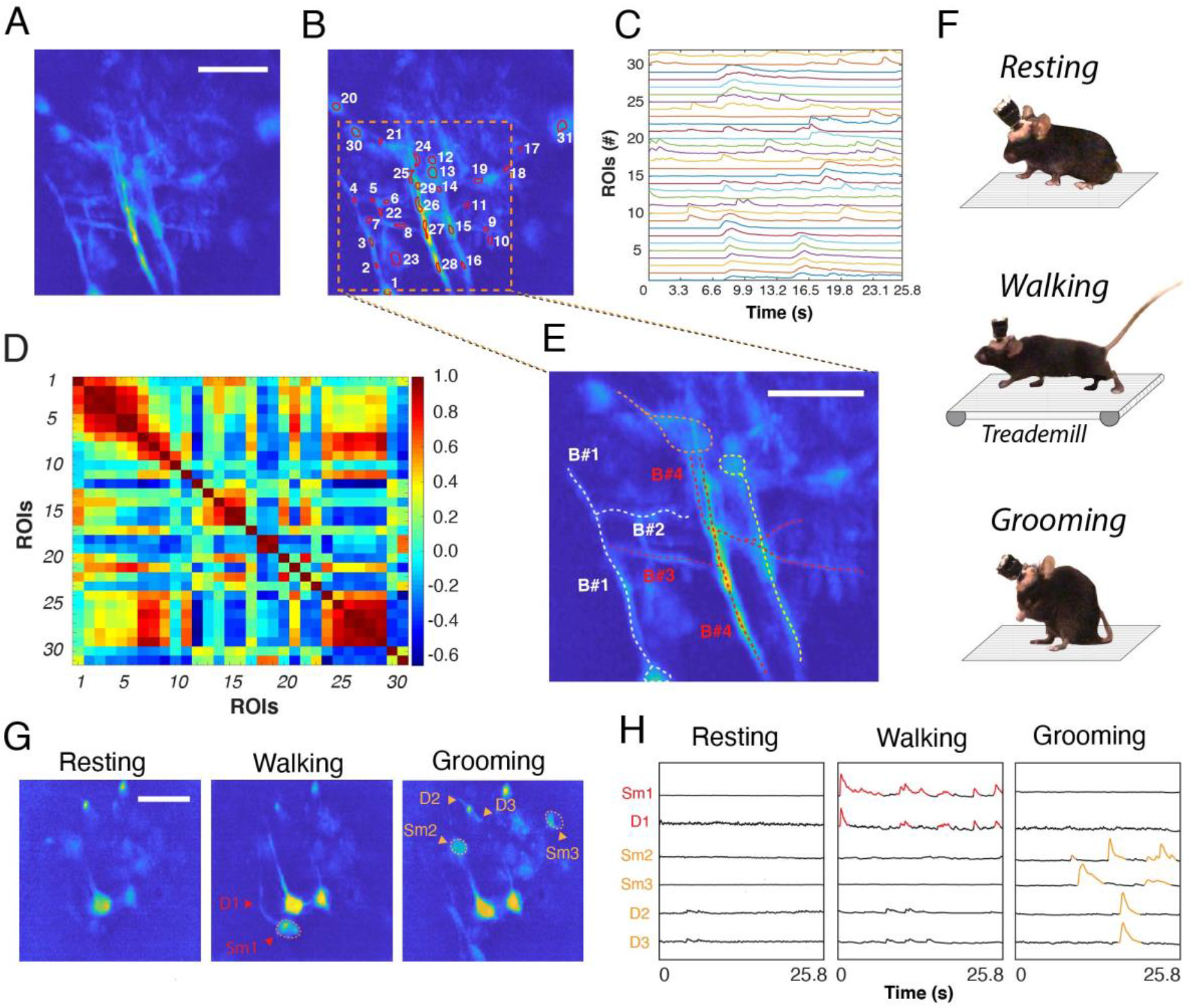
Time-lapse imaging of Ca^2+^ activity from GCaMP6s-labeled neurons of the motor cortex. **(A-D)** Neuronal enhanced calcium activation image **(A),** manually selected ROIs **(B),** calcium traces **(C)** and correlation map **(D)** from the recorded neuronal activity (Figure 4-video supplement 1). Red to dark-blue colors represent maximal (1) and minimal correlation values (−0.6). **(E)** Structural reconstruction based on **D**. Each dendritic branch (B) corresponds to multiple ROIs in **b**; B1: 2,3,4; B2: 5,6; B3: 7,8; B4: 24-29. **(F)** Neuronal activity was imaged and analysed during three behaviours of the animal: resting, walking and grooming. The treadmill was used to encourage the walking comportment while for the remaining conditions, the natural behaviour of the animal was recorded. Neuronal enhanced calcium activation image **(G)** and Ca^2+^ traces **(H)** from somas (Sm) and dendrites (D) during the behaviour in **(F).** Scale bars = 50 μm.

We have then proceeded to the demonstration of the ability of our system to correlate the calcium dynamics of various neuronal structures to the behavioural activity of the animal (Figure 4F). As shown in Figure 4F-H, the level of calcium activity within the entire field of view was constant and equal to the baseline in the animal at rest (Figure 4H). However, the level of calcium activity was modulated in different neurons and their respective fine processes according to the motor activity while the animal was walking or grooming (Figure 4G).

Having confirmed that our system can be used to image and characterize the activity of fine neural processes during different behavioural tasks, we proceeded to the demonstration of the critical advantages enabled by the electrical depth scanning (Figure 5). As shown in Figure 5A, electrical adjustment (55 μm) of the depth position revealed new neuronal features, undetectable at the original working distance of the probe (Figure 5A, top). For instance, within ROI #1 we have discovered a fine dendritic branch (Figure 5A-B, red arrows) that was connected to the pair of newly revealed neurons (orange arrows). A continuation of neuronal prolongation within ROI #2 was also identified (Figure 5C) and quantified (Figure 5D). Thus, the electrical depth shift allowed a better visualization of the neuronal network by enabling the detection of new structures and connections.

**Figure 5.**
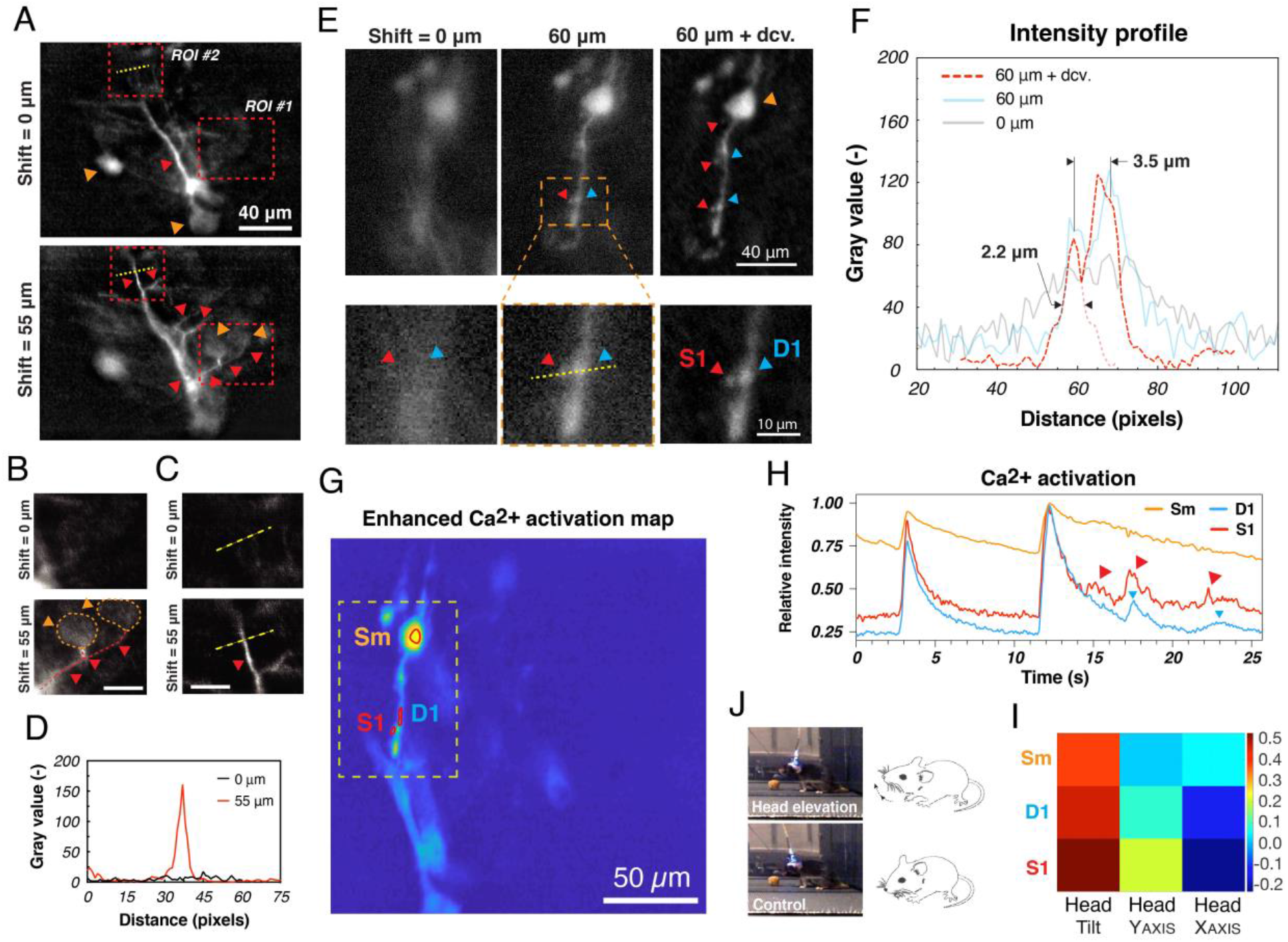
Electrical focus adjustment and time-lapse imaging of Ca^2+^ activity from GCaMP6s-labeled neuronal structures in the motor cortex of a freely behaving animal. **(A)** Images of neuronal activation at different, electrically adjusted focal depths. Focused structures are indicated with arrows (orange: soma, red: processes). **(B-D)** Cropped images **(B,C)** of ROIs 1, 2 and intensity profile **(D)** through the dotted yellow line in **C**. Scale bars = 5 μm**. (E)** Calcium activation images of the soma (Sm), dendrites (D), and spines (S) at two different depths: 0 μm and 60 μm (indicated on the top). Last column image was processed with the Lucy-Richardson deconvolution algorithm (itineration = 15) using DeconvolutionLab2 plug-in (ImageJ). **(F)** Intensity profile through the yellow line in **E**. The diameter of S1 is ≈ 2.2 μm and the distance between S1 and D1 is ≈ 3.5 μm. **(G,H)** Enhanced calcium activation image (MIN1PIPE) with manually selected ROIs **(G)** and the corresponding Ca^2+^ traces **(H)**. Image plane was focused using the TLCL (60 μm). **(I, J)** The head movement **(J)** of the animal was correlated **(I)** to the activity of electrically focused neuronal structures in **G.**

Finally, we have also demonstrated the capacity of our system to electrically focus on even smaller neuronal processes, such as dendritic spines, while examining rhythmic head movements (Figure 5J). The images obtained at the original working distance of the GRIN probe contained only blurry neuronal structures (Figure 5E); electrical adjustment of the working distance (≈ 60 μm) made it possible to clearly identify the soma, dendritic processes and spines (Figure 5F). Furthermore, time-lapse imaging (Figure 5H) showed the capacity of our system to differentiate Ca^2+^ dynamics of a single dendritic spine from its parental dendritic branch and the soma (Figure 5G), which were highly correlated with head movements (Figure 5J, I) of the animal.

## Discussion

In this study, we have presented a miniature depth scanning 1-photon system (mDS1s) that allows multi-plane imaging of neurons and their fine processes, including dendrites and dendritic spines, in freely moving animals. Three fundamental limitations have been successfully addressed.

First, we have developed and incorporated a novel tunable lens (the TLCL) that enabled ≈ 98 μm electrical modulation of the imaging plane within our mDS1s. The utility of this novel feature was shown by imaging neurons (Figure 3E) and fine neuronal processes (Figure 5) that were undetectable in unfocused positions (Figure 5). Electrical modulation also improved the efficiency of surgical procedures by drastically reducing the rate of animal sacrifice due to an incorrect installation depth of the probe. Compared with the commonly used electrowetting lens (Ozbay et al., 2018, 2015), our tunable liquid crystal lens has a 9-times higher optical power (180 diopters) that enables direct optical coupling with small-diameter GRIN probe assemblies (Figure 2D). This allows to achieve a compact optical design (Figure 2) and adaptive imaging with constant magnification (8.7 ± 0.2) during the entire tunability range. The TLCL also has ≈ 20 times lower operation voltages (3 V_RMS_) and ≈ 100-times lower power consumption (≈ μW) compared with electrowetting lens and is, therefore, better suited for its integration within future wireless devices. From a mechanical perspective, the small size (5mm × 5mm × 0.8mm) and the low weight (≈ 0.1 g) of the TLCL are an excellent fit for many miniaturized applications in neuroimaging.

Second, our device shows a high spatial resolution within a 1-photon system capable of imaging fine neuronal processes. To achieve this result, we have designed our device with high magnification along with low optical aberrations (online Methods). Using our mDS1s, we have been able to image, record, and correlate the Ca^2+^ activity of fine neuronal features (e.g., dendrites and dendritic spines) with different behaviors of freely behaving animals. We think that our *in vivo* results will broaden the experimental repertoire of miniaturized 1-photon systems that were so far limited to the imaging of large neuronal structures (Barbera et al., 2016; Cai et al., 2016; Ghosh et al., 2011; Liberti et al., 2017) (e.g., soma). The motionless scanning approach of our mDS1s will highly contribute to the efficient detection and imaging of fine neuronal structures that are particularly hard to detect in absence of perfect focus.

Third, the developed mDS1s also remedies the size and weight limitations of existing depth-scanning devices (Ozbay et al., 2018, 2015; Zong et al., 2017) that can affect animal’s head movement, navigation and multiple decision-making choices. To improve animal’s condition and to preserve its natural behavior, we have developed one of the lightest (≈ 1.4 g) and smallest (10 mm × 17 mm × 12 mm) mD1Ss reported so far (Ghosh et al., 2011). The compact optical design (Figure 2D) and the choice of mechanical frame materials were the key parameters allowing us to achieve this result (online Methods). We think that our mDS1s will provide benefits to the animal and improve the quality of behavioral data in many investigations (e.g., memory, cognition, etc.) based on the natural movement (head and body) of animals.

To summarize, we assert that the advantageous mechanical and optical characteristics of our mDS1s can enable new investigations of fine neuronal structures in physically limited subjects (e.g., younger, older, injured, etc.) that cannot carry currently available heavier devices. Among these subjects are smaller (younger) subjects that are usually used to study neurogenesis and the formation of new dendritic branches and spines of adult-born neurons (Carlén et al., 2002). Older mice that are physically weaker can also benefit from our lightweight device, as one could study the changes in spine density occurring during normal aging (Dickstein et al., 2013) and neurodegenerative pathologies (Bittner et al., 2010; Smith et al., 2009). Finally, our mDS1s might a allow us to assess functional and structural changes in animal models of neurotraumatic injury, such as spinal cord injury (Balbi et al., 2017) or stroke.

In the near future, the development of 2-color, optogenetics-synchronized and wireless versions of our mDS1s should be considered. The low voltage and power requirements of the TLCL will significantly facilitate the engineering of such devices.

## Materials and methods

### Design of the TLCL

We start by building what is called a *half lens #1* sandwich (presented in the bottom of Fig. 1d), by using two glass substrates (100 μm thick). One of the substrates contains a weakly conductive layer (WCL) and a hole patterned electrode (HPE), the combination of both being used to shape in space the electric field and the corresponding alignment of LC molecules. The second substrate is covered by a uniform transparent electrode (indium tin oxide), U-ITO. Homemade nematic LC (NLC) material (presented as filled ellipses) is then filled into the sandwich and aligned (in the plane x,y) almost parallel to substrates by using thin rubbed films of polyimide (PI) on both substrates. This sandwich can be used to focus light with polarization in the x and y plane. Given the non-polarized nature of fluorescence light, a second similar sandwich is then fabricated (see *half lens #2*, top of Fig. 1d) and attached to the first one via a transparent adhesive after rotating it on 90 degrees around the y axis. NLC molecules of the *half lens #2* are thus in the y and z plane and this final assembly can now focus un-polarised light.

### Characterisation of the TLCL

We used a commercial wave-front sensor (WFS150-7AR; Thorlabs) to measure the focal distance and the optical aberrations of the TLCL (Fig. 1e). An electrical square-shaped AC signal was used to activate and drive the lens. Voltage was fixed at optimized value of V_RMS_ = 3.6 V (Figure 1-figure supplement 1) while the frequency of the AC signal was tuned from 1 kHz up to 25 kHz to electrically adjust the optical power (inversely proportional to the focal distance *F*) of the lens.

### Focal shift measurements

Fluorescent beads (FSEG004; Bangs Laboratories, Inc.) were immerged into water and mounted between a standard microscope slice and a thin (4 um) cover glass. Beads were first mechanically defocused using a motorized actuator and then electrically brought back into the focus using the TLCL. The precisely known mechanically out-of-focused distance corresponded to the compensatory electrical shift produced by the TLCL.

### Resolution and PSF measurements

TLCL was used to electrically modulate the working distance of the probe. For each shift value, fluorescent beads were mechanically scanned by steps of 3 μm to generate a 3D point spread function (PSF) stack. The resolution of the system (for each shift value) was determined by using the MetroloJ plug-in (ImageJ) that measured the full width at half maximum (FWHW) of the experimental PSF.

### Animals

All animal experiments were approved by the Animal Welfare Committee of CHU de Québec and Université Laval in accordance with the Canadian Council on Animal Care policy. We used 2-month old male or female VGluT2-IRES-Cre transgenic mice (The Jackson Laboratory, RRID: IMSR_JAX:016963). All mice were either housed alone or with a single sibling after the surgery.

### Surgery

Aseptic surgery was performed under isoflurane anesthesia (1-2%). All pressure points and incision sites were injected subcutaneously with lidocaine-bupivacaine (7.5 mg/kg) for local analgesia. Slow released buprenorphin (1 mg/kg) was administered subcutaneously for post-surgical analgesia. The skull was exposed to do a craniotomy (about 1 mm in diameter) above the motor cortex. The dura mater was removed prior to viral injection. A glass micropipette (ID 0.53 mm, OD, 1.19 mm, WPI) mounted on a nano-injector (Nanoliter 2010 injector with SMARTouch controller, WPI) was filled with AAV2.9/em-CAG-Flex-GCaMP6s (titer 1.2E13 GC/mL) and inserted into the motor cortex to deliver 70 to 100 nL in 2-4 locations within the following stereotaxic coordinates: anteroposterior −0.3 to 0.3 mm from Bregma, lateral 0.8 to 1 mm, depth 0.8 mm. The flow rate was 30 nL/min and after completion, the micropipette was held in place for at least 2 min to prevent leakage. The craniotomy was filled with a hemostat (Bonewax) and the skin above the skull sutured. After a waiting time of at least 3 weeks was allowed a robust expression of the GCaMP6s, mice were instrumented with a GRIN lens during a second aseptic surgery. The implant was secured with self-tapping bone screws (cat#19010-10, FST), crazy glue (exact type) and dental cement (cat# 525000, A-M Systems). Mice were allowed to recover for a least a week before chronic experiments.

### Portability test

To carefully connect the miniscope, animals were anesthetized with isoflurane (2%). Mice were first habituated to the headpiece fixation for 3 days, 30 min per day. Portability test consisted of 4 imaging session spread over two weeks period.

### Data acquisition

At the beginning of each imaging session, headpiece fixation was followed by a resting period of 5 minutes. During each imaging session, the movement of the animal was recorded for 42 minutes at 30 frames-per-second. Imaging session was initiated from software. Mice were allowed to roam free in an open field during imaging without direct contact with the experimenter. Calcium activity in the motor cortex neurons was monitored while recovering from anesthesia, during spontaneous activity such as grooming and during treadmill locomotion.

### Data analysis

Images were analysed using MATLAB and ImageJ. In MATLAB, we used Miniscope 1-Photon Based Calcium Imaging Signal Extraction Pipeline (Min1pipe). The image processing with MIN1PIPE involved motion correction (if required), manual ROI selection and calcium signal extraction. Correlation matrixes were done in MATLAB. ImageJ was used for background subtraction and brightness adjustment. *DeconvolutionLab2* plug-in was used for the deconvolution based on the experimental PSF stacks.

## Supporting information

Time-lapse imaging of Ca2+ activity from GCaMP6s-labeled neurons of the motor cortex.

## Acknowledgements

We would like to thank LensVector *Inc*. for its material and financial support of this work. We are grateful to Dr. M. Andrews and Dr. M. Lemieux for their scientific advices.

## Additional information

### Author contributions

A.B. conceived, designed, built and characterized the system under the supervision of T.G.; A.B., L.T., A.R., and N.K. performed surgery, experiments, and kinematic analysis under the supervision of F.B.; A.B. analyzed the data; A.B., T.G., and F.B. wrote the paper. All authors participated in discussion and data interpretation.

### Ethics

All of the animals were handled according to approved institutional animal care and protocol (#19-027-1) of University Laval. All surgery was performed under anesthesia, and every effort was made to minimize suffering.

## Figure supplements

**Figure 1-figure supplement 1.**
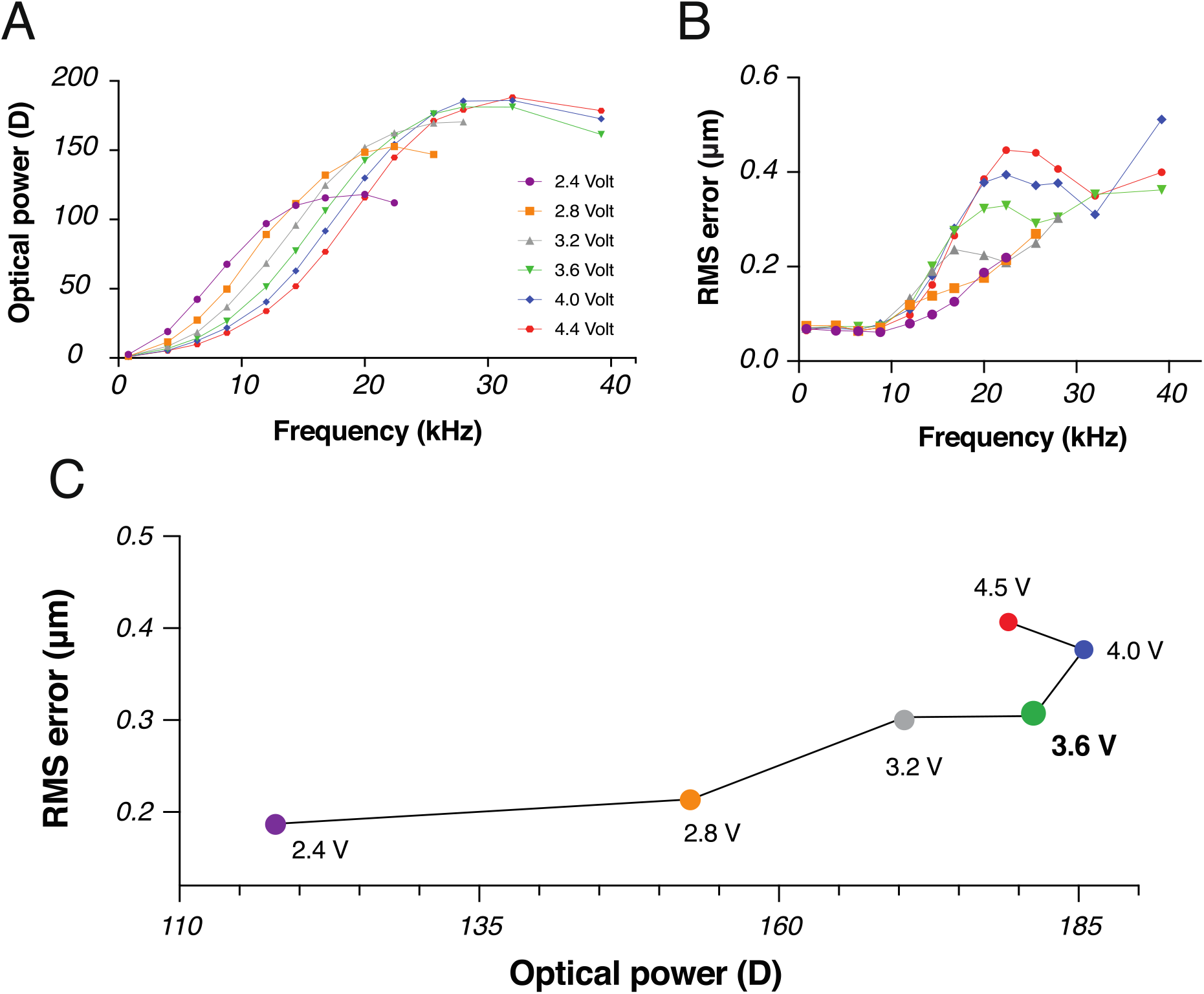
Characterization of the TLCL. **(A,B)** Optical power **(A)** and aberrations **(B)** increase proportionally to the driven frequency of the AC square signal applied to the TLCL. The amplitude of the voltage varied from 2.4 V to 4.4 V. **(C)** RMS aberrations at the maximal optical powers for each of the driven voltages. The optimal voltage: 3.6 V. Voltages bellow that value generated lower optical powers while the higher voltages, had higher RMS error.

**Figure 2-figure supplement 1.**
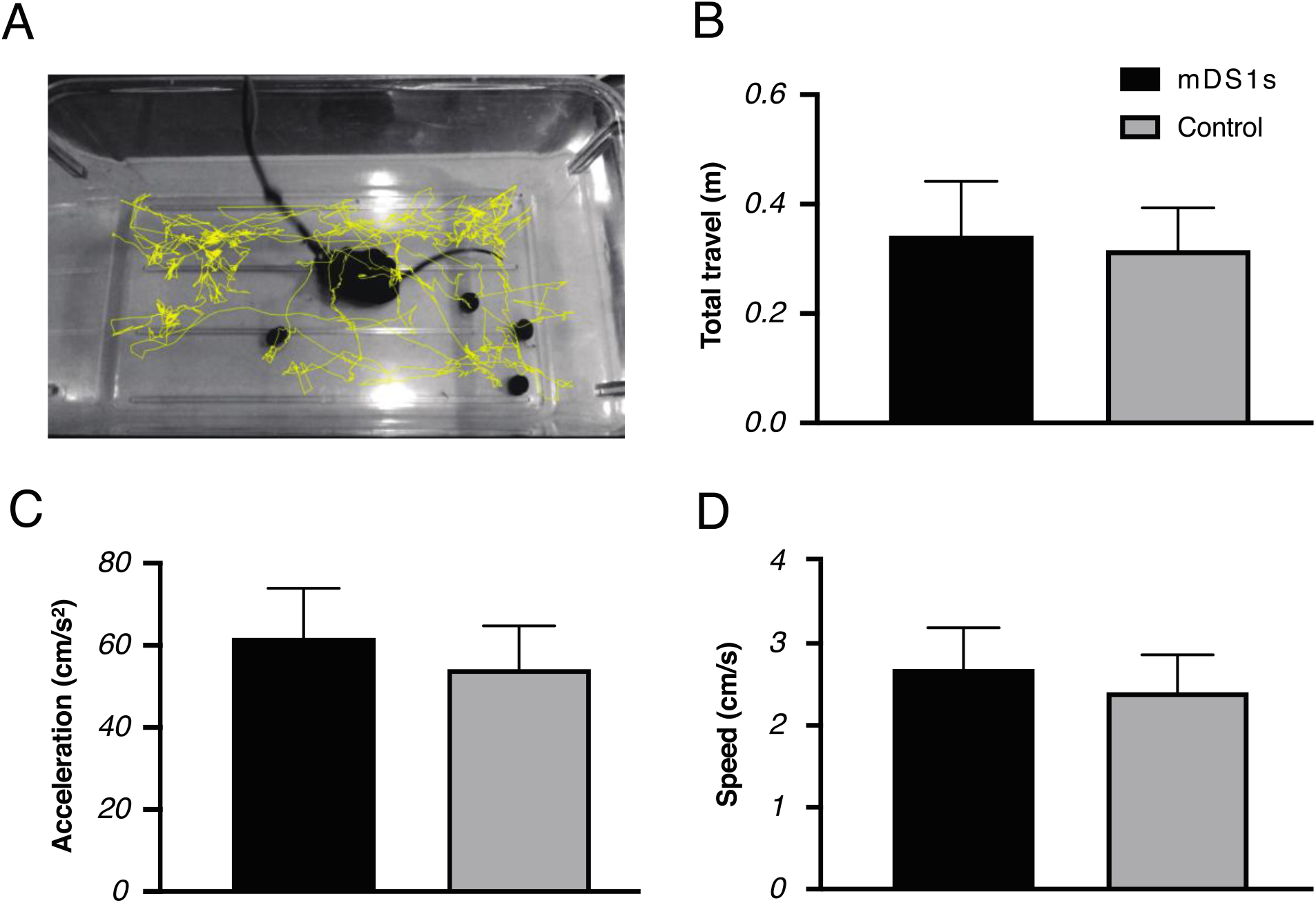
Portability test of the mDS1s. **(A)** Displacement path of the animal during portability test (≈ 4 min). AnimalTracker plug-in (ImageJ) was used to track and analyze animal’s movement. **(B-D)** The total travel **(B)**, the average moving speed **(C)** and acceleration **(D)** of the animal with and without mDS1s. Error bars correspond to the standard deviation after 4 imaging sessions of the animal. Error bars correspond to the standard deviation after 4 imaging sessions of the animal.

